# A drug combination therapy consisting of toxin phospholipase A2 and metalloproteinase inhibitors provides preclinical protection against North American *Crotalid* snakebite envenoming

**DOI:** 10.1101/2025.04.25.648899

**Authors:** Charlotte A. Dawson, Amy E. Marriott, Edouard Crittenden, Adam Westhorpe, Emma Stars, Rebecca J. Edge, Steven R. Hall, Stefanie K. Menzies, Rachel H. Clare, Nicholas R. Casewell

**Author notes:** authors contributed equally (Charlotte A. Dawson, Amy E. Marriott).

## Abstract

**Background:** Across North America an estimated 3,800–6,500 snakebite envenomings occur annually, resulting in 7–15 deaths and an unknown number of disfigurements and disabilities. Most bites are caused by Crotalid snake species. The variable diversity and toxin complexity of crotalid venoms presents a considerable challenge to developing broadly effective small molecule therapeutics to better treat snakebite in this region.

**Methods:** We evaluated the ability of three small molecule, toxin inhibiting, repurposed drugs to inhibit the venom activities of six medically important crotalid snake species (*Agkistrodon contortrix, Crotalus atrox, C. adamanteus, C. horridus, C. scutulatus* and *Sistrurus miliarius*). These drugs target two pathologically relevant venom toxin families, the snake venom metalloproteinases (SVMPs; marimastat and DMPS) and phospholipases A2 (PLA_2_s; varespladib), and venom inhibition was measured using *in vitro* enzymatic and phenotypic plasma coagulation assays. Thereafter we evaluated the efficacy of individual drugs and dual drug combinations in *in vivo* preclinical models of snakebite envenoming, using both preincubation and rescue model formats.

**Results:** *In vitro* bioassays demonstrated that the selected small molecules showed potent inhibition of the enzymatic activity of different toxin families to the nanomolar (varespladib vs PLA_2_ and marimastat vs SVMP) or micromolar (DMPS vs SVMP) level. Three of the venoms had anticoagulant activity, which varespladib restored to normal coagulation profiles, suggesting this activity is mostly driven by PLA_2_ toxins. Preclinical experiments revealed that pre-incubation of representative venoms with single drugs was insufficient to completely protect against lethality, except for varespladib against *C. scutulatus*. Superior efficacy was observed when drugs were used in a combination approach, with the combination of marimastat and varespladib providing greatest protection against lethality in both pre-incubation and rescue models.

**Conclusions:** Venom variation among snake species makes the development of generic snakebite therapeutics challenging. In this study we showed that while SVMP and PLA_2_ inhibiting drugs show inhibitory potency against diverse North American snake venoms, drug combinations consisting of an SVMP inhibitor together with a PLA_2_ inhibitor are required to confer broad *in vivo* protection against lethality caused by envenoming. This study highlights the potential long-term value of drug combinations as next-generation therapeutics for snakebite envenoming.

## 1.0 Background

Snakebite envenoming is a potentially life-threatening event that results from defensive bites by venomous snakes. In the United States of America, 25 species of snakes are categorised as medically important (Gold *et al*., 2005). Although the incidence of snakebite within this region is significantly lower than in the tropics, an estimated 3,800 – 6,500 envenomings and 7 – 15 deaths occur annually in North America (Chippaux *et al*., 2017). Snakebite can also result in life-long morbidity, contributing to a loss in quality of life and potential loss of income through incapacity to work (Gutierrez *et al*., 2017), though accurate incidence estimates from this region are not currently available.

Most snakebites in North America (∼99%) are attributed to vipers from the pit viper subfamily *Crotalidae*, which includes the rattlesnakes (*Crotalus* spp.), the pygmy rattlesnakes (*Sistrurus* spp.) and the moccasins (*Agkistrodon* spp.) (Ruha *et* al., 2017). These snakes have a wide distribution across North America, where they occur in a range of ecosystems, occupy diverse prey niches, and are exposed to a range of climates, all of which are hypothesised to be contributing factors to the high toxin complexity observed in their venoms (Holding *et* al., 2021). Pit viper venoms are a potent yet variable mixture of enzymatic and non-enzymatic components, with the most prevalent toxin families in North American species being the snake venom metalloproteinases (SVMPs) (up to 51% of all venom toxins, dependent on species), phospholipases A_2_ (PLA_2_s) (up to 70%), serine proteases (up to 21%) and C-type lectin-like proteins (CLPs) (up to 38%) (Tasoulis and Isbister., 2017). Common envenoming syndromes resulting from venom toxicity include coagulopathy, localised tissue and muscle damage including necrosis, and in certain cases, neurotoxicity (Chippaux *et* al., 2017, Lavonas *et al*., 2009, Bosen *et* al., 2000,).

Two of the most pathologically important venom toxins families in Crotalids, and therefore the priority for neutralisation following snakebite, are the SVMPs and PLA_2_s. SVMPs are zinc-dependent endoproteolytic toxins that have a broad range of targets within the extracellular matrix (ECM), basement membrane (BM) and coagulation cascade (Dawson *et al*., 2021). For example, SVMPs from the western diamondback rattlesnake *Crotalus atrox* have been shown to target the ECM that surrounds muscle cells, causing extensive tissue damage (Williams *et al*., 2018), while the disruptive action of SVMPs on targets in the coagulation cascade, such as fibrinogen, Factor X and/or prothrombin, can disrupt haemostasis and lead to coagulopathy (Seneci *et al*., 2021, Pandya *et al*., 1983). PLA_2_s have a broad spectrum of activities, with their core enzymatic function relating to the breakdown of phospholipid bilayers (Gutierrez and Lomonte., 2013). This targeted breakdown can contribute towards myotoxicity and oedema in snakebite patients, while biological molecules produced by the breakdown of phospholipids can trigger pathways involved in pain perception and the inflammatory response (Moreira *et al*., 2021, Bickler., 2020; Ferraz *et al*., 2019, Teixeira *et al*., 2003). Perhaps the most pathologically important PLA_2_s found within *Crotalidae* venoms are the neurotoxic PLA_2_s, such as Mojave toxin. Mojave toxin binds irreversibly to pre-synaptic Ca^2+^ channels at the motor-endplate, preventing the binding of the neurotransmitter 3H-nitrendipine (Valdes *et al*., 1989), resulting in fasciculations and paralysis, which can lead to death through respiratory failure (Clark *et al*., 1997). In human envenoming, the presence of these neurotoxic PLA_2_ toxins increases the probability of patients requiring intubation or dying (Massey *et al*., 2012), though they are only found in abundance in the venom of specific populations of certain rattlesnake species (e.g. *Crotalus scutulatus*, *Crotalus helleri*).

Current snakebite treatments, known as antivenoms, consist of animal-derived polyclonal antibodies raised via a process of animal immunisation. Treatment options in the United States and Canada are limited to two antivenoms approved by the U.S. Food and Drug Administration (FDA) – CroFab™ and Anavip™; neither are currently approved by Health Canada, although antivenom can be acquired under special access approval (Brandehoff *et al*., 2023, Wilson *et al*., 2022, Curran-Sill and Kroeker., 2018, Ruha *et al*., 2017). CroFab™ was the only antivenom product available between 2007 – 2018 and is raised against venom from *C. atrox, C. adamanteus, C. scutulatus* and *A. piscivorus,* and is marketed for the treatment of pit viper bites across North America (Wilson *et al*., 2022). Anavip™ became available in 2018, initially marketed for rattlesnake (*Crotalus* spp.) envenomings, then later additionally approved for use against *Agkistrodon* spp. (Brandehoff *et al*., 2023). There is a considerable financial cost to snakebite treatment, especially in the privatised healthcare system of the United States of America. Coupled with the expenses associated with hospital admission, supportive therapeutics, and the need for multiple antivenom vials per patient (median number of vials CroFab; 10, Anavip; 18), individuals can be left with substantial medical bills, with inpatient costs reported to be as expensive as $20,503 (Brandehoff *et* al., 2023; Narra *et al*., 2014).

Given that there are several limitations associated with antivenoms, including high costs, weak efficacy, and high incidence of adverse reactions, next generation snakebite therapeutics have been highlighted as a research priority (Gutierrez *et al*., 2025, Clare *et al*., 2021, Williams *et al*., 2019, Knudsen *et al*., 2018; Knudsen and Lausten., 2018). Various therapeutic modalities for snakebite have been explored to date (Torres *et al*., 2025, Chowdhury *et al*., 2021, Albulescu *et al*., 2020a, Albulescu *et al*., 2020b, Lewin *et al*., 2016, Menzies *et al*. 2023, Slagboom *et al*., 2024, Ledsgaar *et al*., 2023, Edge *et al*., 2024), and small molecule drugs targeting the enzymatic venom toxin families (e.g. PLA_2_, SVMPs) show considerable promise. Recently, increased research effort has focused on the discovery of promising drug leads targeting these two toxin families via the establishment of high-throughput screening approaches (Albulescu *et al*., 2024, Clare *et al*., 2024). However, several late lead compounds that are now progressing into clinical trials were discovered via a rational repurposing approach, followed by validation in preclinical animal studies demonstrating efficacy against severe local and/or systemic envenoming (Abouyannis *et al*., 2025, Bartlett *et al*., 2024, Gerado *et al*., 2024, Hall *et al*., 2023, Menzies *et al*., 2022, Chowdhury *et al*., 2021, Albulescu *et al*., 2020a, Albulescu *et al*., 2020b, Layfield *et al*., 2020, Xie *et al*., 2020; Wang *et al*., 2018, Arias *et al*., 2017, Lewin *et al*., 2016). These molecules include the PLA_2_ inhibitor varespladib and the SVMP inhibitors marimastat (a matrix metalloproteinase inhibitor) and DMPS (a metal chelator).

Such molecules could overcome several limitations currently presented by antivenoms, including oral bioavailability, lower treatment costs, improved safety profiles, the removal of cold chain storage (Clare *et al*., 2021), and the potential for cross-snake species venom neutralisation, particularly in the context of recently described drug combination therapies (Albulescu *et al*., 2020; Hall *et al*., 2024). Whilst several of these attributes seem highly likely to be beneficial for use in low and middle income (LMIC) settings, they could also be advantageous for the North American market. Furthermore, the uptake of next generation snakebite therapeutics in high income countries could be leveraged to support a tiered pricing approach, required to offset the cost of these drugs in LMIC settings, thereby ensuring access to those who suffer the greatest burden of snakebite (Steves and Huys., 2017, Danzen and Towse., 2003).

To assess the suitability of small molecule therapeutics as effective venom-neutralising therapeutics for the North American region, we tested the three aforementioned late lead snakebite drugs (i.e. the SVMP inhibitors marimastat and DMPS and the PLA_2_ inhibitor varespladib) in a range of *in vitro* and *in vivo* venom neutralisation assays against a panel of six North American viper venoms. Here we show that while each inhibitor shows *in vitro* neutralising activity against different pathologically relevant toxins, a drug combination approach is required to provide breadth of neutralising efficacy sufficient to prevent venom-induced lethality caused by different snake species *in vivo*.

## 2.0. Methodology

### 2.1 Venoms

Lyophilised venoms were sourced from the historical venom repository within the herpetarium of the Centre for Snakebite Research and Interventions, at the Liverpool School of Tropical Medicine. Lyophilised pooled venoms from *Agkistrodon contortrix, Crotalus atrox, C. horridus, C. adamanteus, C. scutulatus,* and *Sistrurus miliarius* were stored long term at 2 – 8 °C. Prior to use, all venoms were reconstituted to a stock concentration of 10 mg/mL using PBS (Gibco, pH 7.4, cat no. 10010023) and stored short term at –20 °C.

### 2.2. Small Molecule Drugs

Marimastat (≥98% HPLC, Sigma Aldrich, cat no. M2699-5MG) and varespladib (≥98% HPLC, Sigma Aldritch, cat no. SML1100-5MG) were sourced from Sigma Aldrich and resuspended to 10 mM in DMSO (Sigma Aldrich, cat no: D2650). DMPS (2,3-dimercapto-1-propanesulfonic acid sodium salt monohydrate, 95%, Thermo Fisher, Cat no: H56578) was sourced from Alfa Aesar and resuspended to a concentration of 32 mM in DMSO.

### 2.3. Enzymatic Assays

Established *in vitro*, enzymatic assays measuring SVMP and PLA_2_ activity and plasma coagulation were conducted to determine the neutralising capabilities of the small molecules of interest against the selected venoms (Albulescu *et al*., 2024, Clare *et al*., 2024). Prior to combination with inhibitors, the intrinsic activities of each venom in each of these assays was measured and statistically compared. The AUC of each of the venoms in these assays were compared via one-way ANOVA with Tukey’s multiple comparisons test, with normality confirmed by Shapiro-Wilks test.

#### 2.3.1 Inhibition of snake venom metalloproteinase activity

SVMP activity was quantified by measuring the cleavage of a fluorogenic substrate (R&D BioSystems, cat no. ES010) in the presence or absence of inhibitors. SVMP-inhibiting drugs were serially diluted 1:2 (marimastat 10 μM to 1.22 nM, and DMPS 16 mM to 312.5 nM) and plated at 0.91 µL per well. Each dilution point was assayed in duplicate, with each plate run in triplicate. The PLA_2_-inhibiting drug varespladib (at 1 mM) and vehicle control DMSO were used as assay controls. Venoms were diluted to a working concentration of 0.066 mg/mL and 15 µL added per well to a 384-well plate (Greiner, 781101) to give 1 µg venom/well. Venom and drug were pre-incubated at 37 °C for 25 minutes followed by a 5-minute period of acclimatisation to room temperature. Following incubation, 75 µL of buffer (150 mM NaCl, 50 mM Tris-HCl pH 7.5) containing 1.46 µL of fluorogenic substrate (supplied as a 6.2 mM stock) per 1 mL was added, giving a final assay volume of 91 µL and a final substrate concentration of 7.5 µM. Kinetic fluorescence data was collected using a BMG ClarioStar at an excitation wavelength of 320 nm and emission of 420 nm at 25 °C for 1 hour. Following kinetic data analysis, a single timepoint was selected for each venom, at which activity of the venom-only control had plateaued. From this timepoint, the positive and negative control values were determined, and the inhibitory activity of the compounds at each dose was normalised to percentage of inhibition compared to the negative, no-venom control. From this data, half maximal inhibitory concentration (IC_50_) values were calculated via fitting a nonlinear regression curve to the data points (GraphPad Prism v10.2.3). The IC_50_s of DMPS and marimastat were statistically analysed within each venom to determine significant differences in dose required for inhibition by unpaired two-tailed T test.

#### 2.3.2. Inhibition of venom phospholipase A_2_ activity

Venom PLA_2_ activity was determined using a modified protocol of the commercially available secretory PLA_2_ assay kit (Abcam, cat no. ab133089) (Albulescu *et al*., 2024). First, 10 µL of each venom was plated out into a 384-well plate at concentration ranging from 5 – 0.04 ng/µL. Next, 5 µL of 4 mM DTNB and 30 µL of kit substrate (1.33 mM) were added sequentially to the plate using a VIAFLO automatic pipetting head. The plate was then read kinetically on a Clariostar plate reader at 405 nm at 11 flashes per well with a 161 second cycle time for 15 minutes. Specific PLA_2_ activity (μmol/min/mL) was calculated using the following equation:

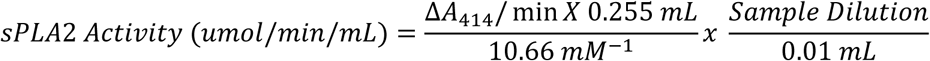

Venom concentrations to be used in downstream inhibition assays were selected based upon the highest dose of venom that did not cause an absorbance plateau during the 15-minute read, indicating that the substrate was not consumed prematurely due to excess venom activity. To evaluate the activity of the PLA_2_ inhibitor varespladib against each of the six venoms, 10 µL of each venom at the selected venom dose (*A. contortrix*, *C. adamanteus*, *C. horridus* and *C. scutulatus* – all 50 ng/well, *S. miliarius* – 30 ng/well, *C. atrox* – 20 ng/well) was incubated with 0.5 µL (10 µM – 1 x 10^-8^ nM, 1 in 10 dilution series) varespladib in the dark at 37 °C for 25 minutes. Plates were then acclimated to room temperature for 5 minutes and read on a ClarioStar plate reader at 405 nm, as described above. The SVMP-inhibiting drug marimastat (at 1 mM) and DMSO were used as assay controls. The rate of substrate conversion was calculated for both the positive (venom-only) and negative (PBS) controls and used to normalise all data to a percentage of the negative control, with 100% representing complete inhibition of venom. From this data, IC_50_ values could be calculated via fitting a nonlinear regression curve to the data points (Graphpad Prism v10.2.3).

#### 2.3.3 Inhibition of coagulopathic venom activity

The coagulopathic activity of the selected venoms was measured via a previously described bovine plasma clotting assay (Albulescu *et al*., 2020). Inhibitors (DMPS: 300 µM – 73.2 nM; marimastat 10µM – 2.44 nM; varespladib: 10 µM-0.01 nM) were spotted into each well of a 384-well plate as a 0.5 µL droplet. Inhibitors were then incubated with 10 µl of venom at a concentration of 10 ng/µL at 37 °C for 25 minutes and followed by a 5-minute acclimatisation to room temperature. Following incubation, 20 µL of 20 mM calcium chloride (Sigma, cat no. C1016-500G) was added per well, followed by 20 µL citrated bovine plasma (VWR, cat no. S0260). Prior to use, the thawed citrated bovine plasma was centrifuged at 3000 RCF for 5 minutes and supernatant collected to remove any cellular debris. Plates were immediately read using a BMG ClarioStar at wavelength 595 nm for 1 hour at 25 °C. Following kinetic reads, the kinetic data was fitted with a nonlinear regression curve from which the ½ time to maximum clot formation could be calculated (the time at which the curve reached half its maximum absorbance). This was performed for the positive and negative controls, and each test well absorbance value was normalised to be a percentage of the positive and negative control values; i.e. 100% would be a ½Vmax identical to the no-venom negative control, indicating complete inhibition of the venom components that affect clotting. Venom activities were analysed by one-way ANOVA with Dunnett’s multiple comparisons test for comparison between venoms, with normality confirmed for each venom dataset by Shapiro-Wilks test.

### 2.4. Pre-clinical Efficacy

#### 2.4.1 Animal ethics

Animal studies were conducted under protocols approved by the Animal Welfare and Ethical Review Boards of the Liverpool School of Tropical Medicine and University of Liverpool under project licence PPL No. P5846F90 approved by the UK Home Office in accordance with the UK (Scientific Procedures) Act 1986.

#### 2.4.2. Animal Maintenance

Male CD-1 mice (18-20 g) were purchased from Janvier (France) or Charles River (UK) and acclimatised for 7 days prior to experimentation in a specific pathogen-free facility. Mice were housed in groups of five in Tecniplast GM500 cages (floor area 501cm^2^) containing 120g Lignocel Select wood fibre bedding (LBS Biotechnology, UK), Z-next biodegradable paper-based material (IPS Product Supplies, UK) for nesting, and environmental enrichment (red house, clear polycarbonate tunnel and loft). Room conditions were set at approximately 20-24 °C at 45-65% humidity, 12/12-hour light cycles (400 lux). Mice were allowed ad lib access to irradiated PicoLab 5R53 food (Lab Diet, USA) and reverse osmosis water via an automatic watering system.

#### 2.4.3. Preclinical efficacy via preincubation model of envenoming

Mice were 25-28 g at the start of experiments. No randomisation was used to allocate experimental groups – mice were allocated into cages of five prior to the experiment, with each cage forming one experimental unit. To determine therapeutic efficacy, a version of the WHO-recommended neutralisation of lethality assay, using preincubated venom plus therapy, was employed. Median lethal doses (LD_50_) for *A. contortix*, *C. atrox*, *C. scutulatus*, and *S. miliarius* were obtained from the existing literature (Consroe *et al*., 1992) and 2.5 x LD_50_ was used as the challenge dose for each venom (*C. atrox;* 3.79 mg/kg, *C. scutulatus*; 0.17 mg/kg, *A. contortrix*; 4.99 mg/kg, *S. miliarius*; 4.87 mg/kg) (Supplementary table 1). Mice initially received a 10 mg/kg subcutaneous dose of morphine as an analgesic, 15 minutes prior to venom injections. Experimental groups (1 cage of n = 5 mice) received an intravenous injection of either: (a) venom only (2.5 x LD_50_), (b) venom and a single drug (60 µg) or (c) venom with a combination of varespladib and either marimastat or DMPS (60 µg of each drug). Treatment groups were not blinded. A total of 30 mice were used across all groups, per venom assayed (total animals used for preincubation studies = 120). Marimastat and DMPS were prepared to 1 mg/mL in sterile water and PBS, respectively. Varespladib was resuspended to 5 mg/mL in DMSO (2.5% DMSO in final dose); all doses containing this drug were sonicated (SoniPrep150 plus) for 10 seconds at 10,000 Hz prior to pre-incubation. All experimental doses were prepared to 200 µL in sterile PBS and incubated for 30 minutes at 37 °C before being delivered intravenously via the tail vein using a BD Micro Fine+ 0.5mL insulin syringe and 29G needle (MediSave, cat no. 324824). After dosing, animals were returned to their home cages for observations for 6 hours (end of experimental time course). Euthanasia was performed using rising concentrations of CO_2_ upon observations of pre-defined humane endpoints (e.g. seizure, hind-limb paralysis, haemorrhage, respiratory distress, immobility coupled with loss of righting reflex). Deaths, time of death and survivors were recorded, where time of death/deaths represent implementation of euthanasia at humane endpoints. Kaplan-Meier curves were plotted in Graphpad Prism v10.2.3 using survival data.

#### 2.4.3. Preclinical efficacy via rescue model of envenoming

To better represent a human bite scenario, we employed a second *in vivo* model where mice received treatment after venom challenge. For these experiments the best performing small molecule combinations identified in the pre-incubation assay were used. Mice (experimental units of n = 5 animals) were challenged with an intraperitoneal dose of venom, before immediately receiving drug combinations in a second intraperitoneal injection. Intraperitoneal LD_50_ doses from the literature (Consroe *et al*., 1992) were used to inform the challenge doses used for study, with our goal to identify a venom challenge dose that ensured an ‘all death’ group within an adequate time window to assess therapeutic efficacy (i.e. venom only controls resulting in death within ∼2-4 hours). Resulting intraperitoneal challenge doses were 2.5 – 7 x IP LD_50_ values described in the literature (*C. atrox;* 5.71 mg/kg; *C. scutulatus*, 0.24 mg/kg; *A. contortrix*, 10.5 mg/kg; *S. miliarius*, 6.84 mg/kg) (Supplementary Table 1). These were dosed via intraperitoneal injection (100 µL volume) to groups of five mice, following pretreatment with analgesic morphine, as described above. For experimental groups receiving treatment, combinations of varespladib (120 µg) with either marimastat (120 µg) or DMPS (120 µg) were then immediately administered intraperitoneally (100 µL) into the same side of the peritoneal cavity to which the venom was injected. Drug doses were scaled up to 120 µg/mouse in line with previous studies (Albulescu *et al*., 2020a, Albulescu *et al*., 2020b). Animals were then monitored for the same pre-defined humane endpoints outlined above for 24 hours.

To establish whether full drug bioavailability, achieved through intravenous administration, would offer a greater level of protection than intraperitoneal drug delivery, the ‘rescue’ model was also applied using intraperitoneal venom challenge followed immediately by intravenous administration (tail vein) of varespladib combined with either marimastat or DMPS. Groups of five mice received the same venom and drug doses outlined above, with both doses delivered in a 100 µL volume. Pre-dosing with morphine, animal monitoring (24 hours) and application of humane endpoints were as described above.

## 3. Results

### 3.1 Small molecule drugs inhibit the *in vitro* SVMP and PLA_2_ activities of North American crotalid venoms

Six medically important crotalid species (*A. contortrix, C. atrox, C. adamanteus, C. horridus, C. scutulatus* and *S. miliarius*) were selected for this work, owing to their diverse venom compositions and geographical distributions. Three small molecules were selected, namely marimastat (a peptidomimetic matrix metalloproteinase inhibitor), DMPS (a metal chelator) and varespladib (a PLA_2_ inhibitor), based on their previously described ability to inhibit PLA_2_s and SVMPs, two of the most pathologically important snake venom toxin classes in these species.

SVMP activity was measured via a fluorometric assay and reported in relative fluorescence units (RFU) (Clare *et al*., 2024). Baseline SVMP activity was highly variable, with *C. atrox* and *S. miliarius* having the greatest activity (346,860 ± 6471 and 349,125 ± 9498, respectively), followed by *C. horridus* (262,862 ± 7,384), *A. contortrix* (235,015 ± 6,285), *C. adamanteus* (212,060 ± 6,518) and *C. scutulatus* (36,055 ± 1,597) (Fig 1a). Apart from *C. atrox* compared to *S. miliarius,* which was insignificant (p = >0.05), statistically significant differences between the AUC of each venom were observed (p = <0.0001). IC_50_ curves were performed to compare the ability of marimastat and DMPS to inhibit SVMP activity (Fig 1b). Marimastat IC_50_s ranged from 17.21 to 111.2 nM, whilst IC_50_s for DMPS ranged from 1.4 to 7.3 µM (Table 1). An IC_50_ for marimastat against *C. scutulatus* venom was not calculated due to all doses tested inhibiting the activity of the venom by over 50%, including the lowest dose of 1.22 nM, a result which is likely due to low SVMP activity of this venom coupled with the high inhibitory potency of marimastat. In all venoms except *C. scutulatus*, which could not be compared due to the aforementioned lack of a calculable IC_50_, the IC_50_ dose of marimastat was significantly lower than DMPS (*C. atrox* p = 0.0003; all other venoms p = <0.0001).The difference in IC_50_s between marimastat and DMPS are consistent with previously reported potency differences between these drugs, which reflect their different mechanisms of action (Clare *et al*., 2024, Menzies *et al*., 2022, Albulescu *et al*., 2020).

**Figure 1.**
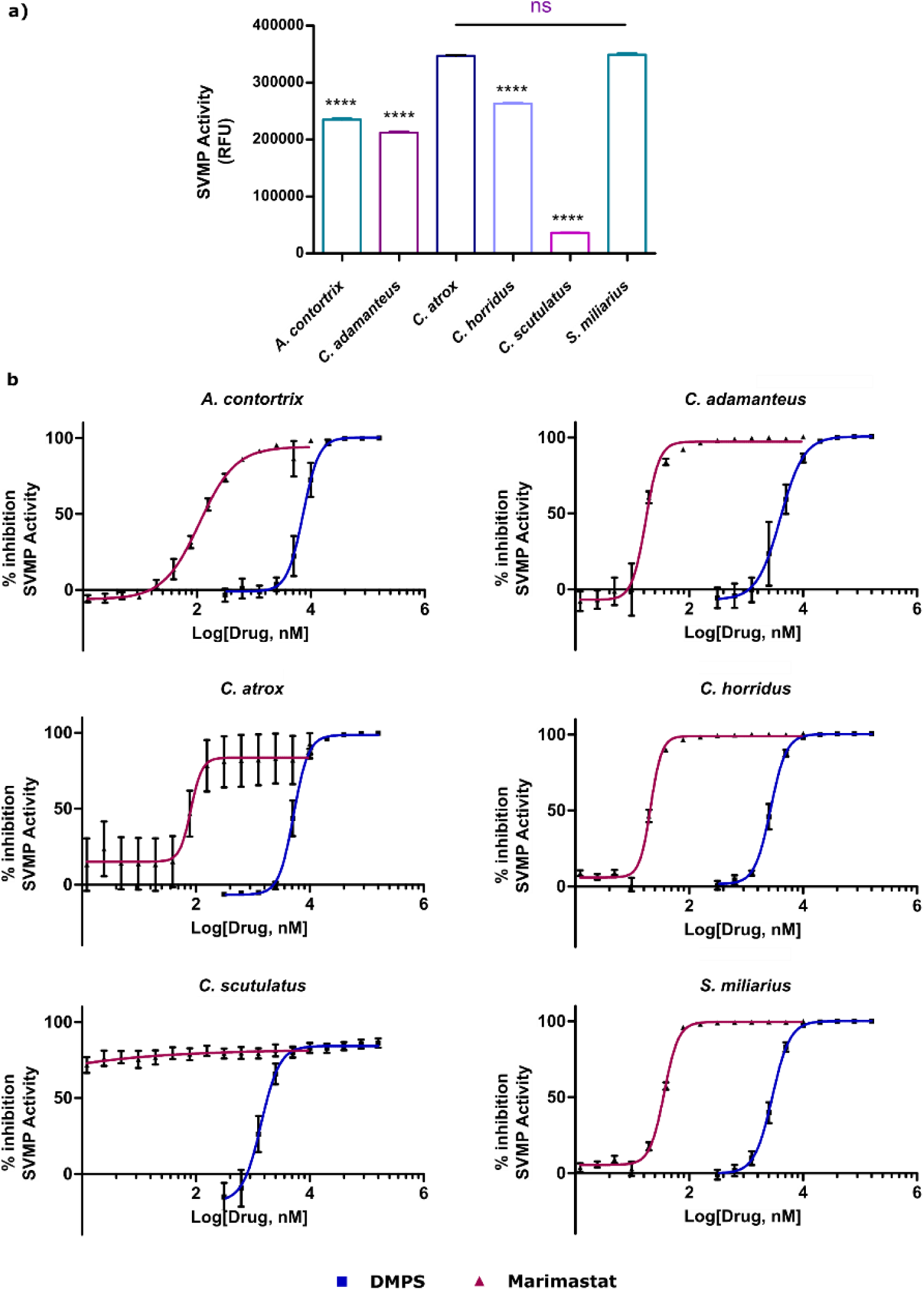
North American crotalids show variable levels of SVMP activity, which is inhibited by DMPS and marimastat. To measure inhibition of metalloproteinase activity by the selected venoms, an *in vitro* fluorogenic assay was applied. a) Enzymatic SVMP activity of 1 μg of venom was measured via cleavage of a fluorogenic substrate (320 nm ex, 420 nm em) at a species-dependent timepoint, shown as average of triplicate reads ± SD. Significance was determined via one-way ANOVA with Tukey’s multiple comparisons test, with normality confirmed by Shapiro-Wilks test. Pairwise comparison between *C. horridus* and *C. scutulatus* was non-significant (ns). All other pairwise comparisons were significant (****) where P = < 0.0001. b) Inhibition of SVMP activity by the metalloproteinase inhibitors DMPS and marimastat. Venoms were pre-incubated with a dilution series of marimastat (10 µM – 1.22 nM) or DMPS (160 µM – 312.5 nM) for 30 minutes at 37 °C before the addition of fluorogenic substrate. Each dilution point was assayed in triplicate, and plotted ± SD. Inhibition of SVMP activity was calculated as percentage inhibition relative to a venom only positive control and no-venom negative control and is plotted against a logarithmic scale of drug concentration, using nonlinear regression (variable slope – four parameters) to produce IC_50_ curves in Graphpad Prism v10.2.3.

**Table 1.** Drug IC_50_s for each venom tested determined in the *in vitro* SVMP (marimastat and DMPS) and PLA_2_ (varespladib) enzymatic assays.

|  | *A. contortrix* | *C. adamanteus* | *C. atrox* | *C. horridus* | *C. scutulatus* | *S. miliarius* |
| --- | --- | --- | --- | --- | --- | --- |
| **Marimastat** | 111.2 nM | 17.21 nM | 79.15 nM | 21.07 nM | < 1.22 nM | 36.21 nM |
| **DMPS** | 7.31 µM | 4.00 µM | 5.20 µM | 2.68 µM | 1.40 µM | 2.87 µM |
| **Varespladib** | 30 pM | 0.50 pM | 0.20 pM | 1 pM | 90 pM | 0.80 pM |

Specific PLA_2_ activity, measured as µmol/min/mL of venom, ranged from 0.52 in *C. horridus* to 4.10 in *S. miliarius* (Fig 2a). Apart from the comparison of *C. atrox* with *C. adamanteus,* which was insignificant (p = >0.05), statistically significant differences between the AUC of each venom were observed (p = <0.0001 – <0.05). Percentage inhibition of enzymatic activity by varespladib was then determined through co-incubation of venom with a dilution series of the drug (10 μM – 10 pM) (Fig 2b). Inhibition assays using varespladib showed strong inhibition of PLA_2_ activity present across all venoms, resulting in a narrow range of sub-nanomolar inhibition from 0.2 pM (*C. atrox*) to 90 pM (*C. scutulatus*) (Table 1).

**Figure 2.**
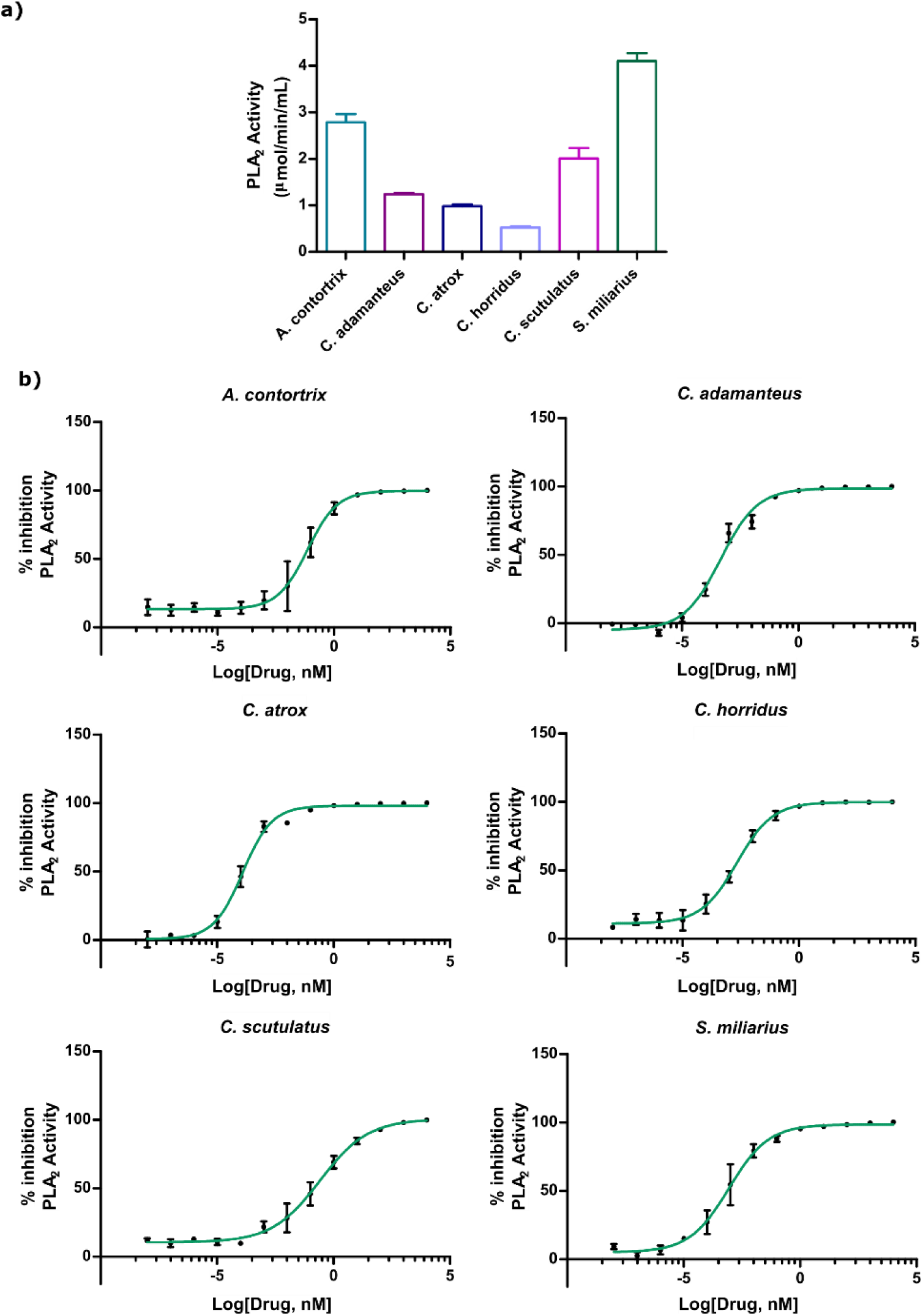
North American crotalids show variable levels of PLA_2_ activity, which is potently inhibited by varespladib. a) PLA_2_ activity (20 or 50 ng venom, depending on species) was measured via colorimetric assay (405 nm) using a kinetic readout over 15 minutes, and enzyme activity calculated as rate of substrate conversion. PLA_2_ activity is presented as a mean of triplicate reads ± SD. b) Inhibition of venom PLA_2_ activity by the small molecule inhibitor varespladib. Venoms were pre-incubated with a dilution series of varespladib (1 mM – 1×10^-8^ nM) for 30 minutes at 37 °C before the addition of DTNB and substrate, with rate of substrate conversion calculated for both positive (venom-only) and negative (PBS) controls used to normalise all data to percentage of the negative control, with 100% meaning complete inhibition of venom. Each dilution point was assayed in triplicate. Percentage inhibition was then plotted (mean ± SD) against a logarithmic scale of varespladib concentration, using a non-linear regression (variable slope – four parameters) to produce IC_50_ curves in Graphpad Prism v10.2.3.

### 3.2 Inhibiting venom activity with varespladib restores normal coagulation profiles

As both SVMP and PLA_2_ toxins can affect the coagulation cascade, we next investigated whether the three drugs could restore normal plasma clotting profiles in the presence of venom. The coagulopathic activity of each venom was first determined using an absorbance-based bovine plasma clotting assay, which revealed that *A. contortrix, C. adamanteus,* and *C. horridus* venoms had no measurable effect on coagulation. The remaining three venoms, from S. *miliarius, C. atrox* and *C. scutulatus,* all exerted anti-coagulant effects on bovine plasma. *C. atrox* and *S. miliarius* displayed significantly different coagulation times (p = <0.05), with no significance seen when comparing any other combination of venoms (p >0.05).

Next, the three anticoagulant venoms were incubated in the presence of the three toxin-inhibiting drugs, and the impact on plasma clotting was quantified by calculating the ‘half time to maximal clotting’ (Fig 3a-c). All three venoms significantly increased the time for clot formation compared to normal clotting (14.66 ± 2.14 mins); *C. atrox* 35.06 ± 2.41 mins (p = <0.0001), *C. scutulatus* 30.33 ± 4.09 mins (p = <0.0001), and *S. miliarius* 23.84 ± 5.89 mins (p = 0.0003). When compared to venom alone, the pre-incubation of *C. atrox* with either SVMP inhibitor resulted in significant reductions in the time to clot formation, with marimastat reducing clotting time to 27.99 ± 3.66 min (p = 0.0035), and DMPS reducing time to 27.37 ± 2.41 mins (p = 0.0017) (Fig 3a). Whilst these reductions in clot time were significant, they remained elevated compared to normal clot formation and no rescue of time to clot formation was seen following the pre-incubation of marimastat or DMPS with *S. miliarius or C. scutulatus* venom (Fig 3b-c). In all venoms, pre-incubation with the PLA_2_ inhibitor varespladib demonstrated a significant reduction in clotting time when compared to venom alone (*C. atrox* and *C. scutulatus*, p = <0.0001; *S. miliarius*, p = 0.0057). Time to clot formation was reduced to within the range of normal clotting when each of the venoms were pre-incubated with varespladib. These findings strongly suggest that the anticoagulant venom effects observed in the species tested here are predominately driven by PLA_2_ toxins, with only a minor contribution from SVMPs.

**Figure 3.**
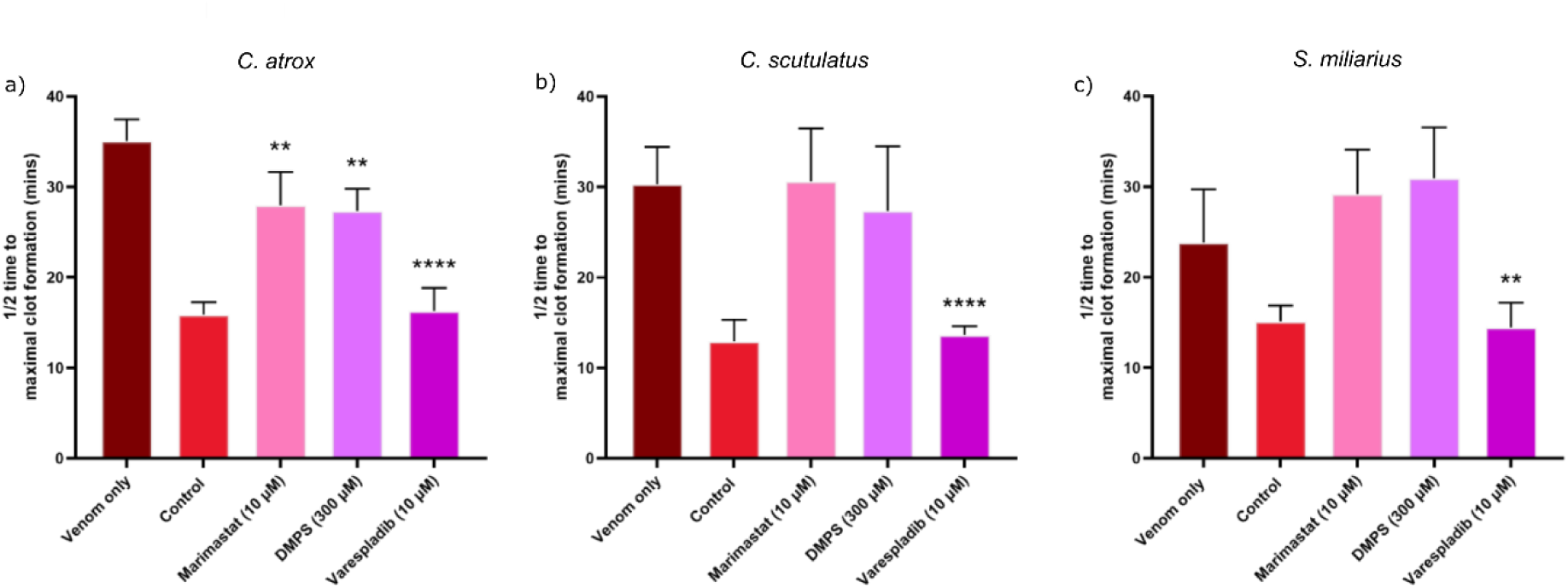
Varespladib is required to restore normal clotting of bovine plasma exposed to North American crotalid venom. Coagulopathic activity of venom was quantified using citrated bovine plasma in the presence of 100 ng of venom and CaCl_2_. Three of the selected North American venoms were found to be anticoagulant; a) *Crotalus atrox,* b) *C. scutulatus* and c) *Sistrurus miliarius.* Application of toxin inhibitors was explored through the preincubation of venom small molecule drugs (30 mins at 37 °C) to inhibit the activity of either SVMPs (marimastat, 10 µM and DMPS, 300 µM) or PLA_2_s (varespladib, 10 µM). Plasma clotting was measured via absorbance assay (595nm, for 1 hour at 25 °C). Clotting in the presence of venom ± toxin inhibitors was presented as the “1/2 time to maximal clot formation”. Statistical significance was measured via one-way ANOVA with a Dunnett’s multiple comparisons test between different drug groups, comparing against the venom only control. ** p = < 0.005, *** P = 0.0009, **** p = < 0.0001. Statistical analysis and graph preparation was completed using Graphpad Prism v10.2.3.

### 3.3 Combination therapies outperform individual drugs in a murine pre-incubation model of snakebite envenoming

We next assessed the preclinical efficacy of the individual drugs in a murine pre-incubation model of snakebite envenoming, where venom and drug were intravenously injected concurrently (Knudsen *et al*., 2020; Menzies *et al*., 2022). We performed these experiments with four of the six venoms we used for *in vitro* assessments (*A. contortrix*, *C. atrox*, *C. scutulatus*, and *S. miliarius*), with those selected based on ensuring coverage of variation in taxonomy, geographical distribution and venom composition. Administration of previously defined 2.5 x LD_50_ doses of *A. contortrix*, *C. atrox,* and *S. miliarius* venoms resulted in rapid murine lethality, with all experimental animals succumbing within 6 minutes, whereas the venom of *C. scutulatus* had a slower onset via the intravenous route, with full lethality occurring at 143 minutes (Fig 4a). This difference was primarily due to the latter venom causing progressive neurotoxicity, rather than the rapid onset haemotoxic effects observed with the other three venoms. Appropriate humane endpoints were selected depending on the observed systemic effects of each venom. For haemotoxic venoms (*A. contortrix, C. atrox,* and *S. miliarius*), observation of seizure, external signs of haemorrhage and laboured breathing constituted a humane end point, whereas the loss of righting reflex, laboured breathing and hind limb paralysis were observed with the neurotoxic *C. scutulatus* venom. For *A. contortrix* and *C. atrox* venom, neither of the SVMP inhibitors (marimastat, DMPS) extended survival times, while treatment with varespladib provided a very modest increase in survival, from a venom only mean survival time of 1.2 and 3 mins compared with 86.3 and 25 mins with varespladib, respectively) (Fig 4a). For *S. miliarius*, marimastat increased survival times in two experimental animals to 272– and 325-minutes post-venom challenge, though the remaining three animals succumbed rapidly to systemic envenoming within 7 mins, in line with the venom only control (complete lethality within 6 mins), while DMPS provided no protective effects at the dose tested (Fig 4a). Varespladib increased survival times compared with the venom only control, with animals surviving between 175 – 342 minutes post-injection (mean of 229.1 mins vs 1 min). Individual inhibitors produced a different profile against the neurotoxic venom of *C. scutulatus*. Co-incubation of venom and varespladib showed strong protection with all experimental animals surviving to the end of experiment (360 mins). Both SVMP inhibitors also provided a degree of protection against this venom, with experimental animals receiving marimastat and DMPS surviving for up to 293 and 316 mins, respectively (mean survival times of 171.2 and 250.8 mins).

**Figure 4.**
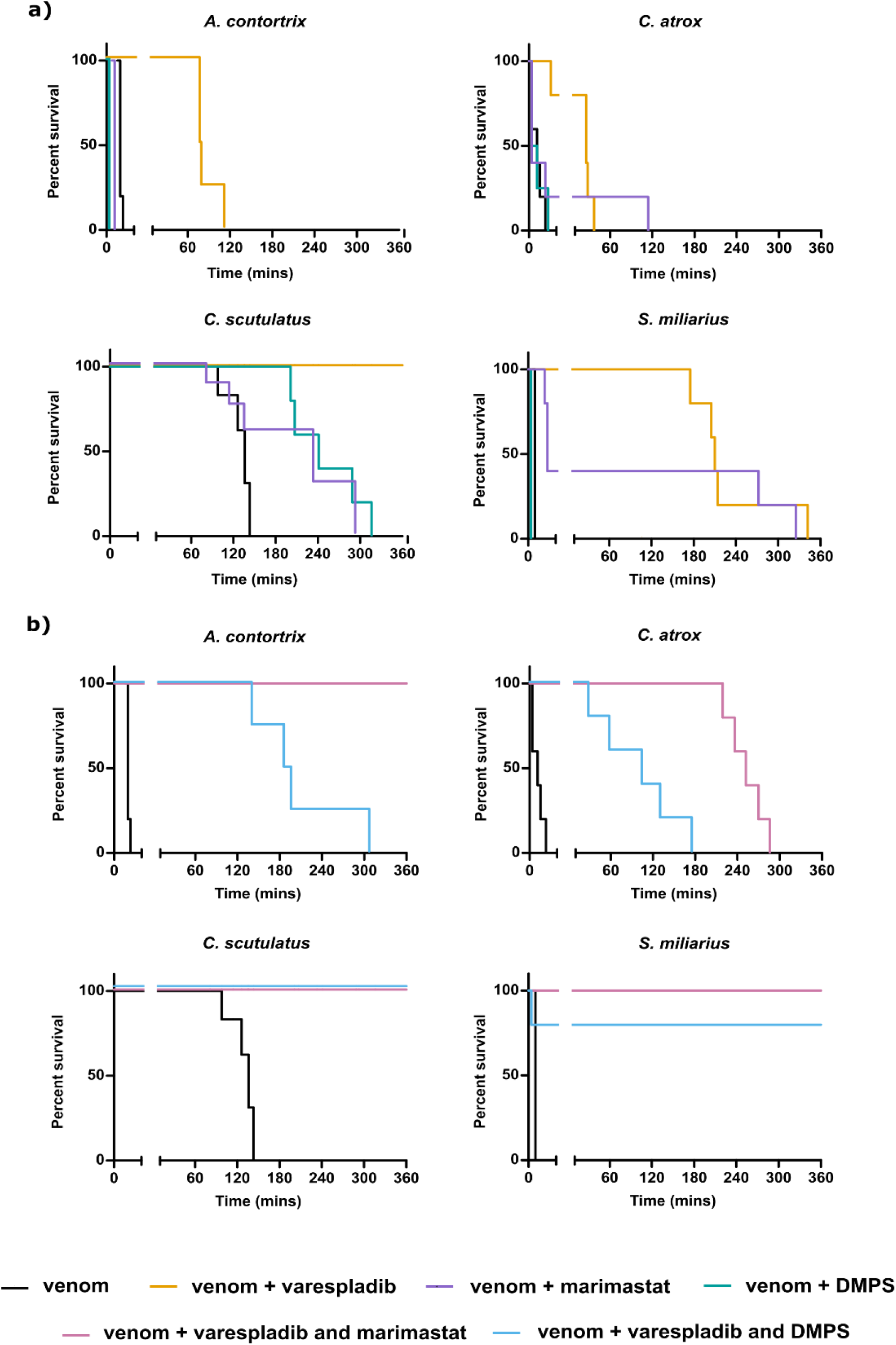
Treatment with a combination of PLA_2_ and SVMP inhibitors provides superior efficacy over single drugs in a preclinical preincubation model of snakebite envenoming. Preclinical efficacy of small molecule inhibitors against selected crotalid venoms was determined via a preincubation model. Venom challenge doses were selected as 2.5 x LD_50_s (*A. Contortrix;* 4.99 mg/kg, *C. atrox;* 3.79 mg/kg, *C. scutulatus;* 0.17 mg/kg, *S. miliariu*s; 4.87 mg/kg). Venoms were then preincubated with either one of the selected small molecule drugs (varespladib, marimastat or DMPS, all at 60 µg) or with a combination of either DMPS and varespladib or marimastat and varespladib (60 µg of each drug in each combination). Groups of n = 5 male CD1 mice were given either: a) venom preincubated with an individual toxin inhibitor or b) venom preincubated with a combination of toxin inhibitors. For each venom, n = 5 mice received a venom alone control. Animals all received an intravenous injection of venom ± therapeutic. Animals were closely monitored for 6-hours for the development of predefined humane endpoints, at which point animals were humanely euthanised via a rising concentration of CO_2_ and time of death recorded and used to generate percentage survival curves in Graphpad Prism v10.2.3.

Use of SVMP and PLA_2_ inhibitors in a combination therapy (varespladib with either marimastat or DMPS) provided superior protection against venom-induced lethality caused by the three haemotoxic venoms (*A. contortrix, C. atrox* and *S. miliarius*) compared to use of the inhibitor monotherapies (Fig 4b). For both *A. contortrix* and *S. miliarius*, the combination of varespladib and marimastat provided complete protection, with all animals surviving to the end of experiment (360 mins). Protection from the varespladib and DMPS combination was also apparent, though more modest, with mean survival times against *A. contortrix* venom extended from 1.2 mins (venom only control) to between 140 – 307 mins, while four of the five experimental animals dosed with *S. miliarius* venom survived the duration of the experimental time course (360 min). Neither combination therapy was able to provide full protection against *C. atrox* venom at the doses tested, though prolonged survival times were observed in mice dosed with both the varespladib and marimastat (252.6 mins, mean survival) and varespladib and DMPS (99 mins, mean survival) treatments, compared with the venom only controls (3 mins). Unsurprisingly, both inhibitor combinations were able to confer full protection against *C. scutulatus* to the end of the experimental time course (360 mins), likely predominately as the result of the inclusion of varespladib, which was fully protective as a solo therapy against this venom.

### 3.4 Combination therapies provide protection against venom lethality in a murine rescue model of envenoming

Based on the superior breadth of efficacy observed when administering both SVMP– and PLA_2_– inhibiting drugs simultaneously in comparison to individually, drug combinations were selected for all subsequent *in vivo* studies. A ‘rescue’ (or ‘challenge then treat’) approach was employed to evaluate the efficacy of the drug combinations when delivered after venom challenge, in a model widely considered to be more reflective of clinical envenoming (and more challenging from an efficacy standpoint) than the traditional preincubation model (Knudsen *et al*., 2020, Albulescu *et al*., 2020, Arias *et al*., 2017). In line with prior studies, we delivered the viper venoms intraperitoneally followed by an immediate dose of the combination therapy in the same location, to enhance the potential to observe efficacy due to local proximity to toxins.

All four venoms (*A. contortrix*, *C. atrox*, *C. scutulatus* and *S. miliarius*) resulted in murine lethality by 117 mins when delivered intraperitoneally. The drug combination consisting of marimastat and varespladib outperformed the DMPS and varespladib combination, in line with the increased *in vitro* and *in vivo* potencies of marimastat over DMPS previously observed (Fig 5a). For envenoming by both *A. contortrix* and *S. miliarius*, the marimastat and varespladib combination provided complete protection over the 24-hour experimental time course against venom-induced lethality, whereas the DMPS and varespladib combination increased survival times against the venom only controls for *S. miliarius* (mean of 667.8 mins [∼11 hrs], vs venom only, 61.2 mins), though only modestly against *A. contortrix* venom (mean of 195.6 mins vs venom only, 60 mins) (Fig. 5a). While the marimastat and varespladib combination therapy also outperformed the DMPS and varespladib treatment against the venoms of *C. atrox* and *C. scutulatus*, the protection provided in this intraperitoneal rescue model was modest, with the former extending mean survival times from 75 and 61.2 mins to 217.4 and 244.6 mins, respectively. The combination of DMPS and varespladib did not provide any protection beyond the venom only control for *C. atrox* (75 mins and 65.5 mins respectively), whilst against *C. scutulatus* the combination of DMPS and varespladib provided modest protection beyond the venom only control (177.7 mins v 61.2 mins, respectively) (Fig. 5a).

**Figure 5.**
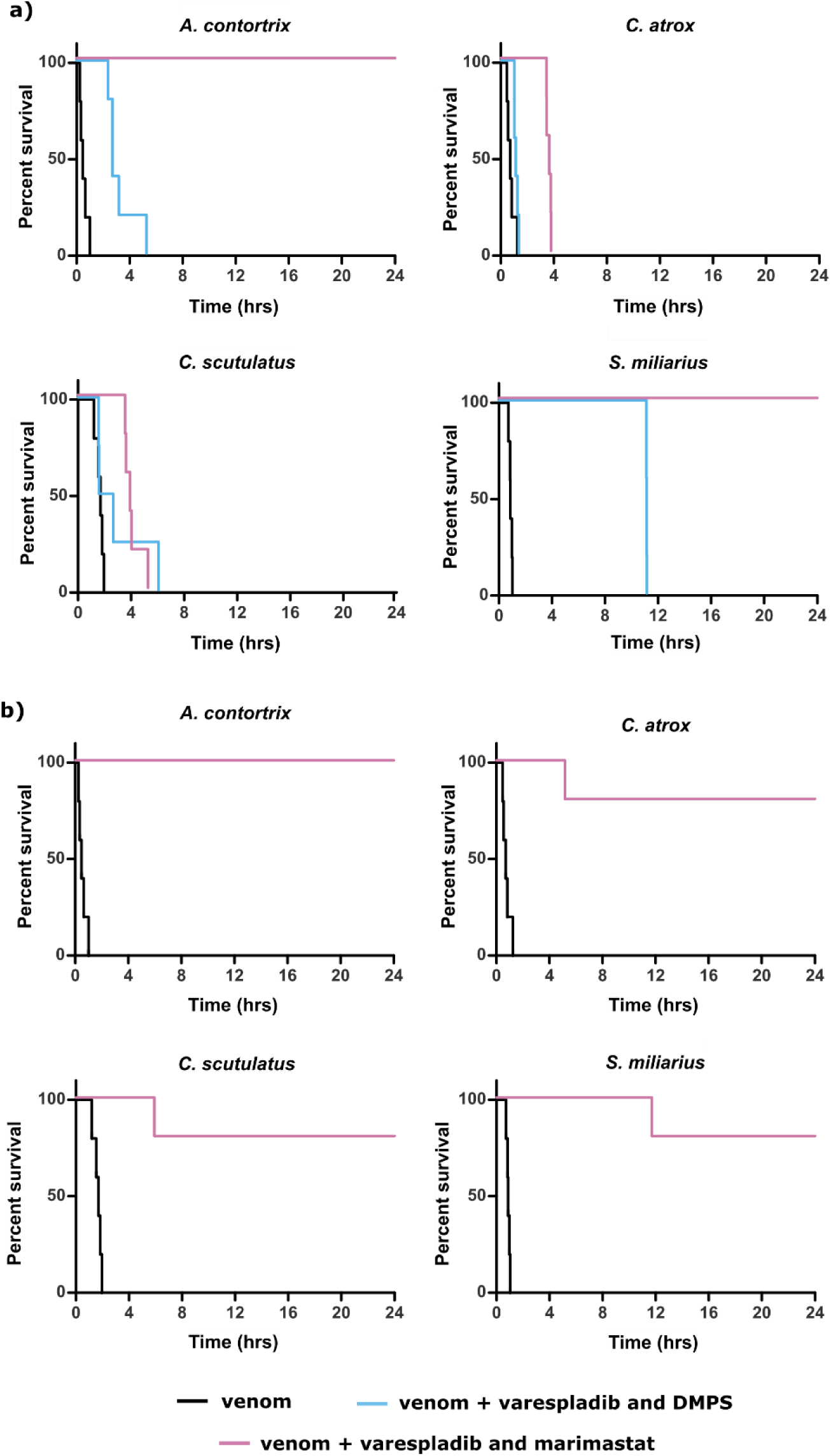
Intravenous delivery of a combination of PLA_2_ and SVMP inhibitors confers protection in a 24-hour preclinical rescue model of snakebite envenoming. Combinational use of small molecule inhibitors was evaluated in a rescue (‘challenge-then-treat’) murine model. Male CD1 mice (n = 5 per group) were challenged with an intraperitoneal dose of venom alone or venom immediately followed by delivery of combined toxin inhibitors. Small molecules combinations consisted of either DMPS and varespladib or marimastat and varespladib (120 µg of each drug). Animals received an intraperitoneal injection (100 µL) of venom at a dosage sufficient to cause lethality within 4 hours (*Agkistrodon contortrix*; 10.5 mg/kg, *Crotalus atrox*; 5.71 mg/kg, *Crotalus scutulatus*; 0.24 mg/kg, *Sistrurus miliarius*; 6.84 mg/kg). Immediately after intreperitoneal venom delivery animals received either: a) an intraperitoneal injection of the drug combinations (100 µL) or b) an intravenous injection of the varespladib and marimastat drug combination (200 µL). Animals were then returned to their home cages and monitored closely for 24-hours. Euthanasia was performed via a rising concentration of CO_2_ upon the observation of predefined humane endpoints. Time of death was recorded upon these observations and used to generate percentage survival curves in Graphpad Prism v10.2.3.

As both drug combinations provided only modest efficacy against the two *Crotalus* spp. venoms in the rescue model, despite evidence of efficacy in the preincubation model, we next repeated the rescue experiments for the four venoms with the optimal drug combination consisting of marimastat and varespladib delivered intravenously rather than intraperitoneally, thereby ensuring maximal systemic bioavailability. This intravenous delivery of the drug combination immediately after intraperitoneal venom dosing resulted in increased efficacy against both the *C. atrox* and *C. scutulatus* venoms, with four of the five envenomed animals surviving until the end of the 24 hours experimental time course for both venoms (Fig. 5b). Intravenous delivery of this drug combination provided complete protection against *A. contortrix* venom, a finding also observed when the therapy was dosed intraperitoneally, though only four of the five animals survived the duration of the experiment with *S. miliarius* venom, compared with complete protection via the intraperitoneal route (Fig. 5b); a difference likely due to variations in the pharmacokinetic profiles of the drugs when administered via different routes (Grogan and Preuss., 2023). Collectively these findings suggest that the drug combination of marimastat and varespladib is broadly effective against an array of North American crotalid envenomings, in both preincubation and rescue models of murine envenoming. However additional work is required to determine optimal dosing regimens via different routes of administration that are sufficient to provide fully protective drug exposures against diverse snakebite envenomings.

## 4.0 Discussion

Envenoming by crotalid species contributes substantially to the snakebite burden of North America. Whilst fatalities are low in this region, particularly when considering the wider global context of snakebite, there is still the potential for morbidity or mortality, especially when bites occur in remote locations (Chippaux *et al*., 2017). Although antivenom treatment is available across the region, the cost of privatised health care in the United States can leave patients with significant medical costs. Given the high cost of accessing treatment, and the potential for envenomings to occur in remote settings, there is a strong case for exploring the utility of small molecule therapeutics. In this study, we selected a subset of medically important crotalid venoms with diverse venom compositions, geographic ranges and envenoming pathologies, and tested the efficacy of three well-characterised orally bioavailable small molecule therapeutics in a range of *in vitro* and *in vivo* assays. To determine the broad applicability of the small molecules against the six selected venoms, we first used established *in vitro* assays. These assays highlighted the variation in venom activity seen across North American pit viper species. Two species, *C. atrox* and *S. miliarius,* had high SVMP activity, whilst *A. contortrix* and *S. miliarius* venoms were enriched in enzymatic PLA_2_ activity. The pathological importance of these toxins informed the selection of marimastat and DMPS to specifically target SVMPs, and varespladib to target PLA_2_s. These repurposed drugs have demonstrated preclinical efficacy against other medically important venoms and have progressed, or are being progressed, into phase I and II clinical trials for snakebite (Abouyannis *et al*., 2025, Gerado *et al*., 2024, Carter *et al*., 2022). *In vitro* IC_50_ testing revealed that of the two SVMP inhibitors, marimastat was a more potent inhibitor compared to DMPS. Furthermore, the IC_50_ values generated (17.21 – 111.2 nM marimastat, 1.4 – 7.3 µM DMPS) are also comparable to IC_50_s reported against other medically important viperid snake venoms (Clare *et al*., 2024).

Whilst enzymatic assays provide valuable insight, the pathophysiological impact of venoms is more frequently due to a combination of toxin activities, therefore we utilised a phenotypic assay measuring coagulopathy with bovine plasma. Despite the complexity of North American pit viper venoms, the observation the varespladib alone was sufficient to rescue clotting response suggest that the coagulopathic effects of these venoms are at least driven by the action of PLA_2_s. The consistent efficacy of small molecules throughout the range of *in vitro* assays applied, and the limitations of studying the complex pathological effects of venom *in vitro*, provided strong rationale for progressing their evaluation into preclinical snakebite models.

Previous studies have shown that venom pre-incubation with individual small molecule drugs can be sufficient to confer full protection against lethality caused by certain venoms *in vivo* (Albulescu *et al*., 2020). However, in this study, when each inhibitor was tested for preclinical efficacy against four crotalid venoms, only varespladib was able to confer full protection against a single venom, *C. scutulatus*. Given that progressive neurotoxicity was observed in animals challenged with *C. scutulatus* venom, which is well established to be driven by the presynaptic PLA_2_ ‘Mojave toxin’ (Gopalakrishnakone *et al*., 1980), the efficacy of varespladib alone against this venom is perhaps unsurprising. Against the remaining venoms, none of the single drug therapies of varespladib, marimastat or DMPS were able to prevent venom lethality from occurring across the full 6-hour experimental time course. This lack of protection seen, even with the optimal model conditions of following venom pre-incubation and co-administration, is likely the result of the relative toxin complexity of North American crotalid venoms (Tasoulis and Isbister, 2017).

Prior preclinical experiments suggest that small molecule therapeutics used in combination may offer broader protection across snake species compared to the use of inhibitors in isolation (Albulescu *et al*., 2020; Hall *et al*., 2023). Evidence of the benefit of combination therapies was observed here with both drug combinations tested conferring greater protection against lethality compared to individual drugs against three of the four pit vipers tested. In preincubation experiments, the combination of marimastat and varespladib conferred greater protection compared to varespladib and DMPS, with complete survival of experimental animals observed for at least 99 mins following lethality in venom only controls. Prior *in vivo* studies have demonstrated that marimastat has greater toxin inhibitory potency than DMPS, likely due to the direct inhibitory mode of actions compared with indirect metal chelation which may explain the efficacy differences observed here (Menzies *et al*., 2022, Chowdhury *et al*., 2022, Jones *et al*., 2022, Albulescu *et al*., 2020).

The combination of marimastat and varespladib also provided greater protection against lethality when animals were dosed via the intraperitoneal route in a rescue model of envenoming. However, complete survival of experimental animals was only observed against two venoms suggesting that reduced drug bioavailability via intraperitoneal dosing may be a limiting factor. When dosing the marimastat and varespladib drug combination via the intravenous route in the same rescue model, we observed prolonged survival of experimental animals (with a minimum of 80% survival) across all four venoms. Collectively these findings suggest that optimisation of drug doses and route of administration requires further exploration to ensure sufficient drug levels are circulating throughout the experimental time course. For example, the dose of DMPS used here, which was dose matched to marimastat, may simply be insufficient for DMPS to deliver a fully protective effect that could potentially be conveyed at higher therapeutic doses or in a multiple dosing regimen. Thus, detailed pharmacokinetic (PK) and pharmacodynamic (PD) analysis of lead candidate molecules in animal models is likely to be highly informative for optimising dosing regimens and informing the value of potential onward translations.

The data generated here provides considerable support towards the use of small molecule drug combinations as novel therapeutics for snakebite envenoming. As the body of preclinical evidence for these drugs expands, clinical data relating to their use as solo therapeutics is now beginning to emerge. Varespladib has entered phase II clinical trials across North America and India. While clinical trial outcome measures were not met in the first of these studies, sub-study analysis showed treatment with varespladib in conjunction with antivenom resulted in a greater percentage of patients showing full recovery by day 28 post-envenoming compared to those in the placebo group (antivenom only) (Gerado *et al*., 2024), while a second trial is ongoing at the time of writing. Additionally, DMPS recently completed a Phase I clinical trial that escalated the oral dose of this already licenced drug for snakebite indication, resulting in a well-tolerated, PK-informed, multiple oral dosing regimen ready for use in Phase II (Abouyannis *et al*., 2025). Given that the primary therapeutic use of these drugs would be as oral medications delivered promptly following a bite and prior to secondary antivenom (i.e. due to barriers associated with access to tertiary healthcare facilities housing antivenom), the efficacy of the drug combinations described herein should next be evaluated preclinically via the oral route, ideally rationally informed by robust PK/PD data. Future clinical trials will require the use of small molecules in combination with an appropriate antivenom, therefore further preclinical experiments modelling the dual approach should also be explored, ideally mimicking early oral drug delivery followed by delayed intravenous antivenom, as previously outlined for DMPS (Albulescu *et* al., 2020). Lastly, the efficacy of these small molecule combinations should be further evaluated in models of localised tissue damage following envenoming using crotalid venoms. Prior preclinical studies have shown that both the marimastat and varespladib and DMPS and varespladib drug combinations significantly reduce the development of dermonecrosis caused by *C. atrox* venom (Hall et al., 2023). Given the extensive variation in North American crotalid venom, this work should be expanded to encompass additional medically important crotalid species to better evaluate the potential impact of these drug combinations on morbidity caused by snakebite.

Whilst the scale of the medical impact of snakebite in North America differs from that of many LMIC settings, there remains a clear need for new therapeutic avenues across the region. Given the documented venom variation between different North American crotalid species, we hypothesised that a drug combination approach of simultaneously targeting SVMP and PLA_2_ toxins with inhibitory molecules would be required to deliver broad cross-snake species preclinical efficacy. Through both *in vitro* and *in vivo* evaluation of three repurposed small molecules, we have been able to demonstrate the value of this drug combination approach. These findings provide both an exciting avenue towards the improvement of snakebite treatment in North America and, perhaps most importantly, open up the potential to leverage this commercially viable market to support the future development and ensure the long-term accessibility of new snakebite therapeutics to LMICs via tiered pricing structures.

## Declarations

### • Ethics approval and consent to participate

All animal experiments were conducted using procedures approved by the Animal Welfare and Ethical Review Boards of the Liverpool School of Tropical Medicine and the University of Liverpool, and in accordance with the UK Animal (Scientific Procedures) Act 1986. All experiments were conducted under UK Home Office approval, in accordance with the conditions of Project Licence P58464F90.

### • Availability of data and materials

All data generated or analysed during this study are included in this published article [and its supplementary information files].

### • Author contributions

CAD, AEM, RHC, and NRC conceptualised the project and designed the experiments. CAD, AEM, AW and RHC performed *in vitro* experiments. CAD, AEM, EC, RJE, SRH, ES, SKM and NRC performed *in vivo* experiments. CAD, AEM, AW and NRC analysed and interpreted resulting data. CAD and AEM wrote the first draft of the manuscript, which was reviewed and edited by AW and NRC. All authors read, edited and approved the submitted manuscript.

## • Acknowledgements

We thank Paul Rowley for the maintenance and husbandry of the snake collection and the provision of venom samples at LSTM. The authors also acknowledge use of the Biomedical Services Unit provided by Liverpool Shared Research Facilities, Faculty of Health and Life Sciences, University of Liverpool.

## List of abbreviations

WHO: World Health Organisation
FDA: Food and Drug Administration
SVMP: Snake Venom Metalloproteinase
PLA_2_: Phospholipase A2
CLPs: C-type lectin-like proteins
LMIC: Low and Middle-Income Country
DMPS: 2,3-dimercaptopropane-1-sulfonic acid

---

**Supplementary Table 1.** Venom doses used in the *in vivo* preclinical studies.

| Venom | Preincubation model |  | Rescue model |  |
| --- | --- | --- | --- | --- |
|  | Intravenous<br>(LD<sub>50</sub> mg/kg) | Intravenous<br>(µg/mouse) | Intraperitoneal<br>(LD<sub>50</sub> mg/kg) | Intraperitoneal<br>(µg/mouse) |
| *Agkistrodon contortrix* | 4.99 | 224.55 | 10.5 | 472.5 |
| *Crotalus atrox* | 3.79 | 170.55 | 5.71 | 257.0 |
| *Crotalus scutulatus* | 0.17 | 7.65 | 0.24 | 30.24 |
| *Sistrurus miliarius* | 4.87 | 219.15 | 6.84 | 430.92 |

